# Rampant Reverse Evolution towards Specialization in Commensal Bacteria Colonizing the Gut

**DOI:** 10.1101/096206

**Authors:** A. Sousa, R. S. Ramiro, J. Barroso-Batista, D. Güleresi, M. Lourenço, I. Gordo

## Abstract

The maintenance of diversity in the gut microbiota is a signature of host health. Yet how strain variation emerges and changes over time in this ecosystem is poorly understood. Here we use a natural yet controlled system to track the effects of natural selection by the genetic signatures it leaves in evolving populations. By following the emergence of intra-species diversity in an *Escherichia coli* strain, we unravel a recurrent case of violation of Dollo’s law, which proposes that evolution is unidirectional and irreversible. We demonstrate *de novo* acquisition of a primordial lost phenotype via compensatory mutation and also genetic reversion, the latter leaving no trace of the past. We show that this reverse evolution generates two coexisting phenotypes, resource generalist and specialist, whose abundance can be controlled by diet supplementation. While specialists’ abundance is low, they avoid competition with the gut microbiota, whereas generalist abundance is dependent on microbiota composition. Our results highlight how a single genetic change can have large ecological consequences.

---

The diversity of bacterial species inhabiting the host intestine is important for its health (1). Furthermore, within each bacterial species, many different strains can colonize the gut but the evolutionary mechanisms by which such intra-species diversity is structured remains elusive (2, 3). In particular the dynamics of emergence and spread of adaptive mutations in commensal gut bacteria is poorly known (4). Recently using the streptomycin treated mouse model of *E. coli* gut colonization, and a fluorescently labeled strain, we began to uncover the nature of the adaptive process that this bacteria experiences in the gut. We found that the first steps of its adaptation, arising as rapidly as three days post-colonization, consisted in the selective inactivation of the operon that allows *E. coli* to metabolize galactitol (5), a sugar alcohol derived from galactose. Distinct, but phenotypically equivalent, knock out alleles of this operon bearing a similar selective effect (7.5% benefit (6)) were recurrently observed to emerge across independent populations adapting to the mouse gut (7). Thus, the distribution of fitness effects of mutations comprising the first step of adaptation to the gut could be described by a simple Dirac-Delta distribution (8). Using this experimental system, we unravel the spread of adaptive mutations, by following the dynamics of two neutral fluorescent markers in an otherwise genetically monomorphic population (5). Here, we have enquired if the pace of adaptation would slow down in the next steps of adaptation to this complex ecosystem. A deceleration in the pace of adaptation is expected under simple models of adaptation (9) and is typically found in bacterial populations adapting to laboratory environments (10–12). If the selective benefit of emerging mutations decreases during adaptive walks, then diversity might be maintained for longer periods of time.

## Results

We colonized 15 mice with a clonal population of *E. coli* exhibiting the first beneficial phenotype – inability to uptake galactitol-conferred by a single base pair insertion in the coding region of the *gatC* gene, which encodes a subunit of the galactitol transporter. The colonizing population was made dimorphic for a neutral fluorescent marker to allow determination of the spread of further adaptive changes (8). The time series dynamics of the fluorescent marker frequency resulting from the second steps of adaptation are shown in Fig. 1a. These dynamics can be summarized in two effective evolutionary parameters (5, 8, 13), the mutation rate (*U_e_*) and the fitness effects of adaptive mutations(*S_e_*) that are estimated to be *U_e_*= 7.1 x 10^−7^, *S_e_*= 0.09 (see Methods). These quantitatively describe the simplest adaptive scenario where new beneficial mutations emerge at a constant rate and have a constant effect (8). The estimates of the evolutionary parameters for the second steps of adaptation are remarkably similar to those previously estimated to cause the first step (*U_e_*= 7 x 10^−7^, *s_e_*= 0.075) (5). This analysis indicates that the pace of adaptation is similar following the spread of the first adaptive phenotype. Direct *in vivo* competitions between samples of evolved clones and their ancestor further corroborate that the fitness benefit of the accumulated mutations is large, ranging from 5 to 14% fitness increase (mean fitness increase of 9% (SD=4%)) (Fig. 1b). Thus, unlike what is typically observed in *in vitro* evolving populations (10–12), the strength of selection *in vivo* does not seem to decrease.

**Fig. 1.**
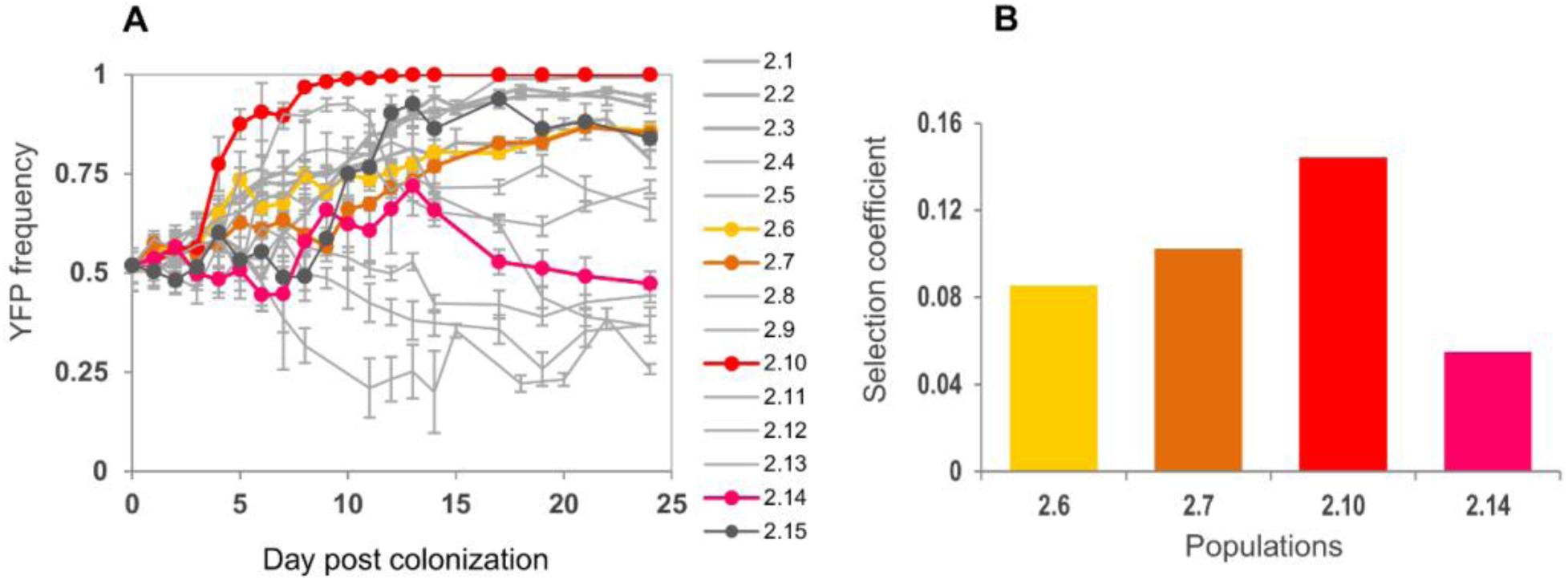
High rate of adaptation characterizes the evolution of *E. coli* colonizing the mouse gut. (**a**) Change in frequency of a neutral fluorescent marker (YFP) along 24 days of colonization. (**b**) Fitness increase (per generation) measured by direct *in vivo* competitive assays of evolved clones against the ancestor (*n*=2).

To determine the genetic basis of adaptation underlying the rapid changes in neutral marker frequency (Fig. 1a) we performed whole genome sequencing (WGS) of samples of two evolving populations along time in two lineages (2.14 and 2.15 from Fig. 1a-b).

Remarkably, we uncovered the rapid emergence of several mutations in the galactiol operon (Fig. 2a-b, see also Table S1), particularly in the previously pseudogenized *gatC* (Fig. 2c). These mutations restored the open reading frame of *gatC* suggesting that an unexpected re-gain of function could have evolved. Thus we tested hundreds of evolved clones from these populations for ability to metabolize galactitol and found that the new mutations in *gatC,* spreading through the populations (Fig. 2a-b), were indeed gain of function mutations. In face of our previous findings (5), where *gat*-positive *E. coli* evolved a gat-negative phenotype that reached more than 99% frequency in half of the mice after 24 days of colonization (Fig. 3a), the re-emergence of the *gat*-positive phenotype was completely unexpected. Indeed we had assumed that the *gat*-negative phenotype would eventually fix in all populations and therefore started a second colonization with a single adaptive *gat*-negative mutant. This design, mimicking a strong bottleneck or transmission event, serendipitously revealed that evolution could reverse.

**Fig. 2.**
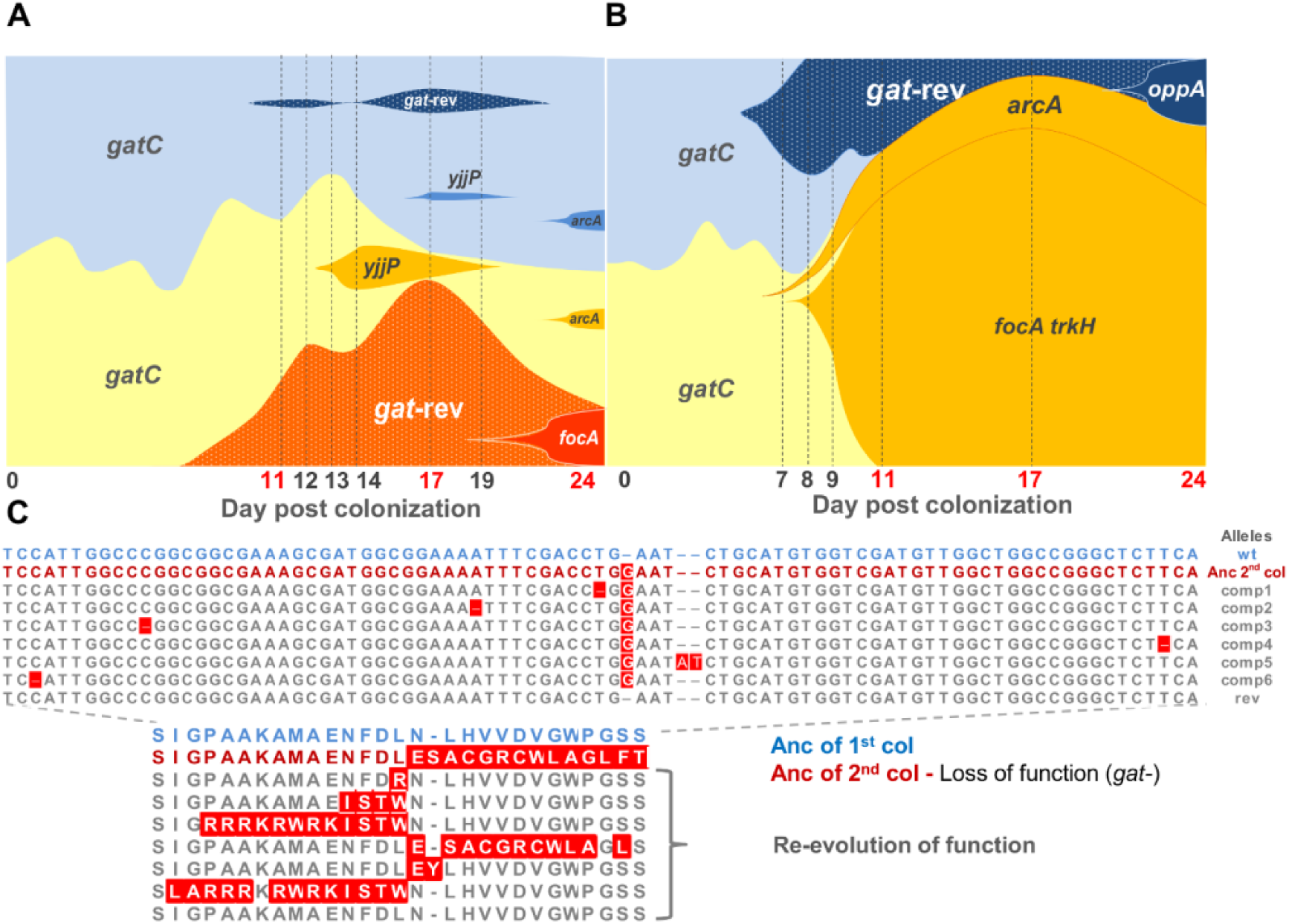
Regain of a previously lost phenotype with recurrent violation of Dollo's law. (**a** and **b**) Muller Plots showing the emergence of polymorphism in two evolving populations (pop 2.14 (**a**) and pop 2.15 (**b**)) where regain of function (ability to metabolize galactitol) is detected - gat-rev. Yellow and blue shaded areas are proportional to the frequencies of YFP- and CFP-labelled bacteria. Darker tones of yellow and blue denote the accumulation of adaptive mutations in the respective background. Days shown below the graph depict the temporal points where haplotype frequencies were estimated (see Table S1) through a combination of different approaches. Numbers in red represent temporal points where frequencies were estimated by WGS of the population and numbers in black represent time points where frequencies were obtained by target PCR of collections of clones. Dotted areas represent the proportion of the population that reverted the ancestral *gat*-negative phenotype and was determined by phenotypic testing. (**C**) The genetic basis of the *gatf-*reversion phenotype (*gat*-rev) involves both compensatory (represented by alleles comp1-6) and back-mutation (allele rev). Six different alleles were found in pop 2.14 at day 11 (comp1 through 6). After 6 days (day17) only alleles comp1 (∼1% frequency) and 2 (∼13% frequency) could be detected by WGS; we note however that the frequency of *gat*-rev phenotype in the same time-point was considerably higher (∼50%). This suggests that the polymorphism in the *gatC* locus estimated by WGS was considerably underestimated in this time-point. On day 24, comp2 was the only allele detected. Days 6 and 11 of pop 2.15 revealed two different alleles (comp3 and rev). On day 17 and 24 only the rev allele was found.

To query about the extent to which the two populations studied reflect a typical outcome for reversion (14), we tested large samples of clones from all the evolved populations for a *gat*-positive phenotype. We found that phenotypic reversion was highly probable, as in 80% of the 15 independently colonized mice the bacteria re-gained the ability to consume galactitol, with this phenotype reaching a detectable frequency of at least 0.3% and up to 77% in the evolving populations (Fig. 3b).

**Fig. 3.**
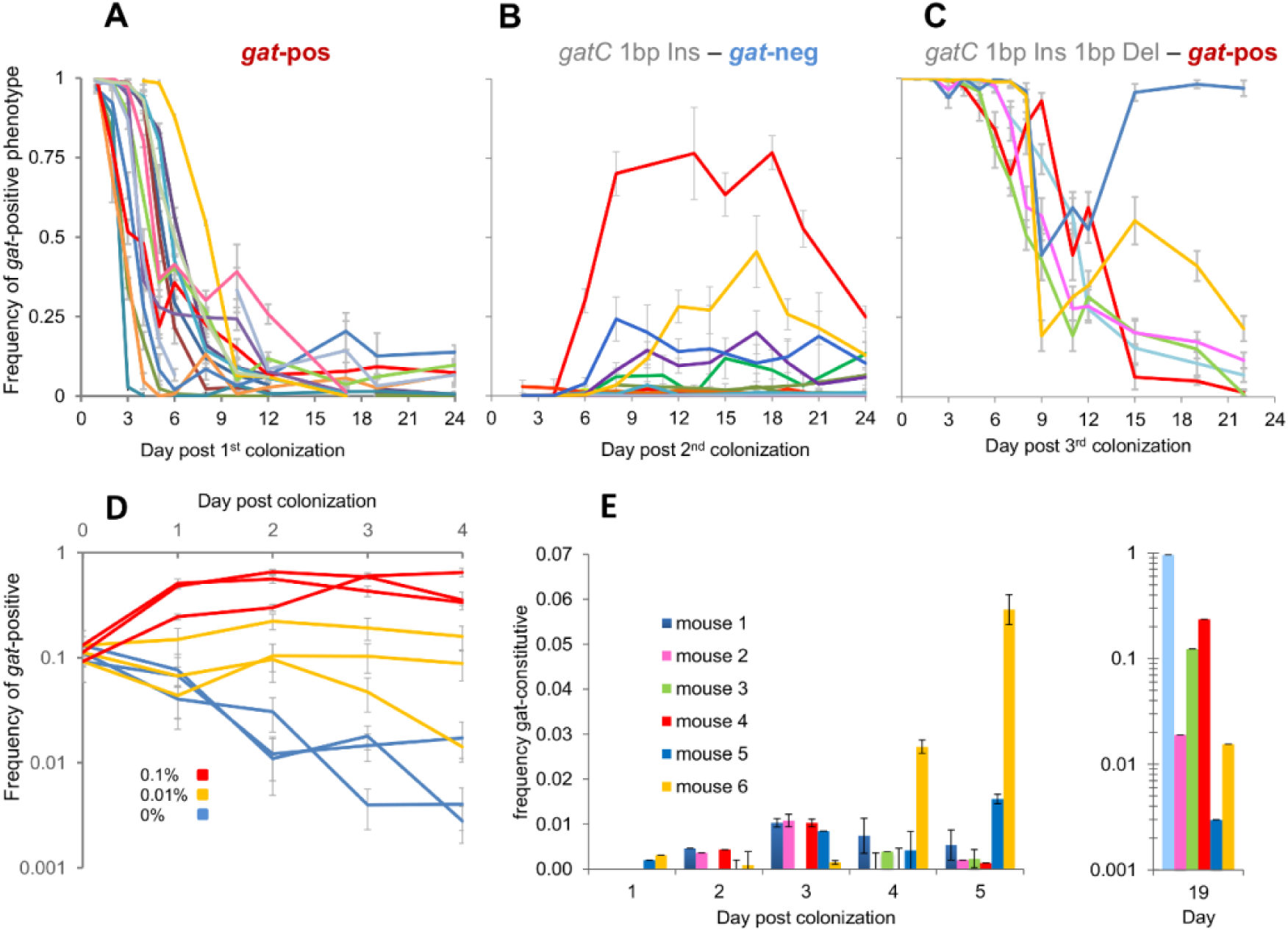
Reacquisition of the ancestral metabolic ability is pervasive across independently evolving populations. (**a-c**) The ancestral genotype and phenotype of each colonization is indicated above the respective graph. (**a**) Massive selection to inactivate the gat-operon characterized the first colonization, yet in half of the populations the frequency of *gat*-positive bacteria was kept above 1%. (**b**) In the second colonization, initiated with a *gat*-negative strain, reacquisition of the ancestral metabolic ability was pervasive across independently evolving populations re-establishing the polymorphism for the *gat* phenotype. (**c**) The third colonization, initiated with a *gat*-positive clone isolated from the second colonization, exhibited the re-emergence of *gat*-negative mutants, as in the first colonization. (**d**) Increase in frequency of the *gat*-positive strain in the guts of mice where diet is supplemented with galactitol. (**e**) Emergence of *gat*-constitutive strains in the guts of mice colonized with a clonal population of bacteria where the repressor of the galactitol operon (*gatR*) has been restored.

In sum, under a transmission event involving a single clone, re-acquisition of polymorphism at the *gat* locus constituted a highly predictable adaptive event. Interestingly, the power of selection on this re-gain of function is clear in population 2.14 (Fig. 2a), where an allele (involving a 1bp deletion, the only mutation detected by WGS as indicated in Table S1) emerges in this population at day 11, on the YFP background, but will then compete with a CFP clone which acquired another mutation causing the same phenotype. Thus the emergence of a *gat*-positive phenotype in both fluorescent backgrounds within the same evolving population (Fig. 2a) and the parallelism across independent replicate populations (Fig. 3b) strongly supports the adaptive nature of the regain of function in this ecosystem (15). To provide further evidence for selection acting on mutations conferring the *gat*-positive phenotype, in contrast to being hitchhiker alleles, we sequenced two *gat*-positive clones. The sequencing of the two clones (isolated from population 2.15 at day 6) revealed that a single mutation had occurred in each clone: a back mutation (“rev allele” in Fig. 2c) and a compensatory mutation (“comp3 allele” in Fig. 2c). These results suggest that the *gat*-positive phenotype is intrinsically beneficial. Moreover, the rare event of gain of function through back-mutation could even be observed in one of the assayed populations (2.15, Fig. 2b-c), supporting the conclusion that the ancestral gene can invade a population where loss of function at this locus had previously evolved. Such an event is particularly important since it erases the signature of the strong past selection that had occurred.

To understand to what extent selection in this ecosystem can overwhelm genetic drift, due to extreme population bottlenecks, in driving the emergence of intra-species diversity at this locus, we performed a third colonization. This was initiated with an evolved *gat*-positive clone (isolated from population 2.14 – Fig. 2a), which carries two additional mutations in the *gatC* gene (see Materials and Methods). In all 6 colonized mice, emergence of the *gat*-negative phenotype, consistent with inactivation of the *gat* operon, was seen (Fig. 3c). In this new genetic background the emergence of polymorphism at detectable frequencies occurred slightly later than in the first colonization: the median time for *gat*-negative phenotype to reach 5% frequency was 7 days (6.25-7.75, Interquartile range), whereas for the first colonization it was 3 days (2-4, Interquartile range, Mann-Whitney-Wilcoxon Test, W=1.5, P=0.0006).

The three colonizations together (Fig. 3a-c) suggest a model of cycling evolution, whereby bacterial populations can loose genetic diversity due to intense drift, associated with transmission/ migration events, but can readily restore polymorphism at this locus due to high mutation rate (*gat*-positive to *gat*-negative 1.0 x 10^−5^ (95% CI, [6.7 x 10^−6^, 4.0 x 10^−4^]) (7) and *gat*-negative to *gat*-positive 9.7 x 10^−9^ (95% CI, [3 x 10^−9^, 1.9 x 10^−8^]) per generation (see Materials and Methods)) and strong selection in this ecosystem.

Competition for limiting resources is a mechanism likely to be important in the gut microbial ecosystem (16). Such a mechanism can lead to strong selection for evolving polymorphisms (17). We hypothesized that galactitol could be a key limiting resource for *E. coli* in the mouse gut and explored the conditions under which a simple resource competition model (18, 19) could explain the invasions observed in the independent colonizations of Fig. 3a-c (see Materials and Methods and Fig. S1). In this simple model we assume that the *gat*-negative and the *gat*-positive clones consume two different resources. Given that the *gat*-positive strain constitutively expresses the *gat* operon (it carries an IS1 insertion in the operon repressor, *gatR*), we further make the simplifying assumption that this strain only consumes one of the resources (galactitol). This simple theoretical model can produce dynamics of emergence of polymorphisms very similar to the observed. The model further predicts that supplementation of the mouse diet with galactitol should lead to an increase in frequency of *gat*-positive clones (Fig. S1.). To test this prediction we colonized new mice with a co-culture of *gat*-positive and *gat*-negative bacteria (at a ratio 1 to 10) while supplementing the mice diet with different concentrations of galactitol. As predicted by the model, the frequency of *gat*-positive bacteria increased in mice drinking water with galactitol (χ^2^_2_=35.8, *p*<0.001), and more so when the concentration of galactitol was higher (Fig. 3d, also compare with Fig. S1b). The model also predicts that a *gat*-positive specialist may be able to invade a resident population of clones that are not specialized at consuming galactitol (see Fig. S2). To query if this *in silico* prediction would be met *in vivo*, we reconstructed a strain with a functional repressor of the galactitol operon (*i.e.* we restored regulation replacing the pseudogene *gatR* by the wild type version) and followed its evolution when colonizing the mouse gut for 19 days (*n*=6). As predicted by the model, and consistent with previous *in vitro* experiments of adaptation to limiting nutrient concentrations (20–22), we observed the emergence of *gat*-constitutive bacteria in all mice (Fig. 3e). Targeted PCR of the evolved clones showed that the phenotype can be caused by *de novo* IS insertions in the coding region of *gatR*. This experiment thus shows that the ancestral clone, which started the first colonization (Fig.3a), can be selected for in the mouse gut, when the mouse is colonized with a strain carrying a functional repressor of the galactitol operon.

Overall the colonization experiments (Fig. 3) imply that irrespectively of the founder clone being constitutive or regulated, in the long run *gat*-negative and *gat*-positive polymorphism is likely to emerge and be maintained in the gut. They also indicate that the selective pressure to evolve clones that specialize on this limiting resource is very high in this environment.

The theoretical model suggests that the equilibrium ratio of *gat*-positive to *gat*-negative is dependent on the relative abundance of galactitol to other undefined resources (see Materials and Methods) and that the abundance of these two phenotypes can vary independently as these consume different resources. Interestingly, in the galactitol supplementation experiment (Fig. 3d), we see that increasing galactitol concentration leads to an increase in the abundance of *gat*-positive bacteria (χ^2^_2_=15.8, *p*<0.001), without affecting the abundance of *gat*-negative bacteria (χ^2^_2_=2.4, *p*>0.05, Fig. S3). Conversely, in the colonization experiment (Fig. 3b) it is the abundance of *gat*-negative bacteria that varies across mice, while the abundance of *gat*-positive bacteria remains relatively constant after their emergence (Fig. 4). This suggests that there is variation in the concentrations of the unidentified resources that are consumed by the generalist, *gat*-negative *E. coli.* This variation could be mediated by the mouse gut microbiota. We thus enquired whether we could observe an association between the loads of *gat*-negative *E. coli* and the microbiota community composition in the gut of the colonized mice (Fig. 4a-b). Remarkably, we find that while all mice show similar loads of *gat*-positive *E. coli* (Fig. 4a-d) mice with a low load of the *gat*-negative *E. coli* (Fig. 4c) indeed exhibit a microbiota community composition significantly different from that observed in mice where high loads of *gat*-negative *E. coli* are attained (Fig. 4d). We observed two main microbiota compositions (UPGMA clusters) that we term cluster I and cluster II (I in red and II in blue; Fig. 4a, c-e). Cluster I shows significantly higher species richness (Shannon index= 4.7±0.2 (2SE); χ^2^_1_=15.5, *p*<0.001; Fig. 4e), being enriched for several OTUs (5 Firmicutes, 2 Bacteriodetes and the Defferibacteres *Mucispirillum;* Fig. S5). Cluster II shows a lower Shannon index of 3.3 (±0.3), being enriched for a single OTU from each of Actinobacteria (Coriobacteriaceae), Proteobacteria (Entobacteriaceae) and Bacteriodetes (*Bacteriodes*).

**Fig. 4.**
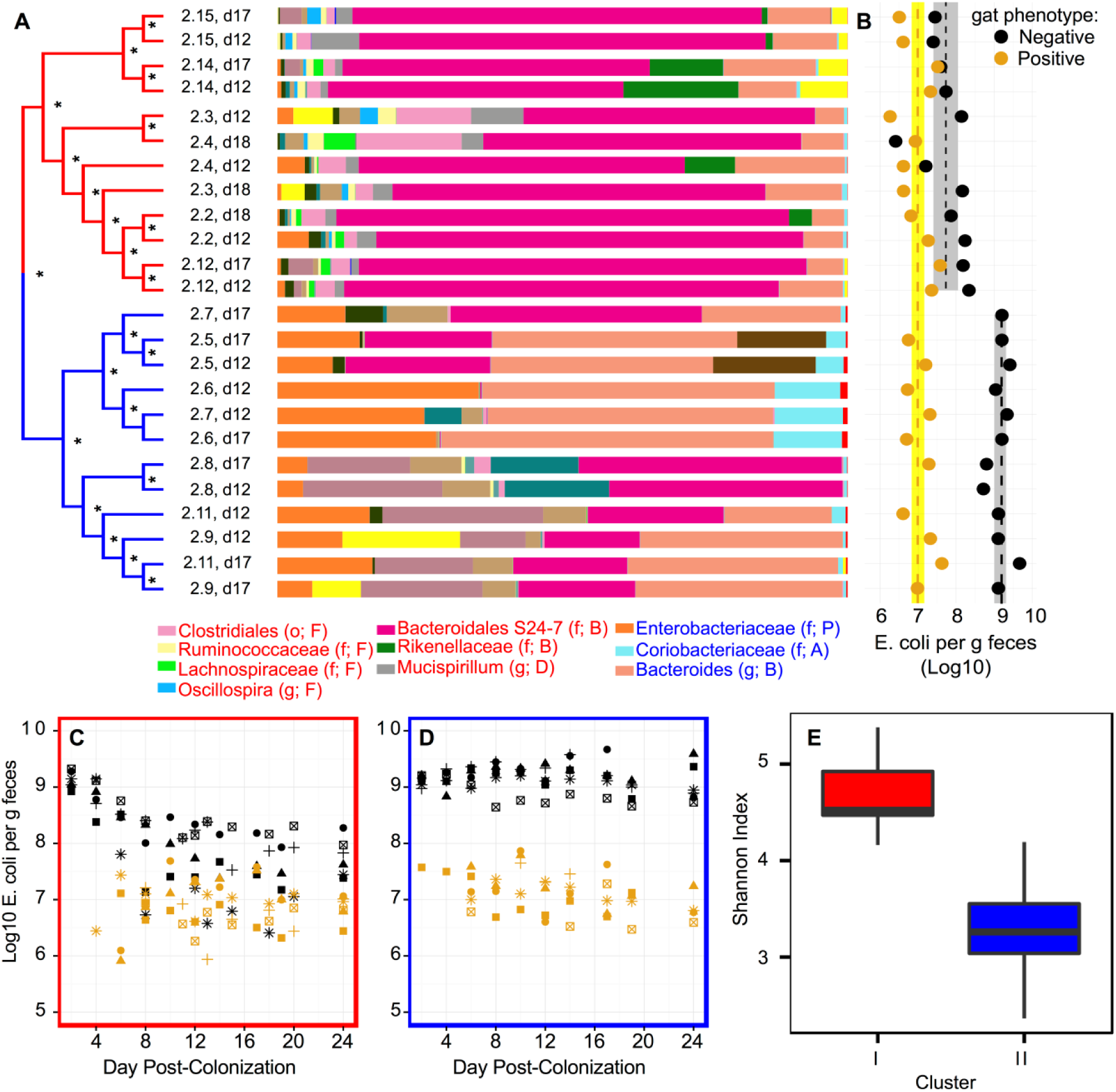
Microbiota composition affects the loads of a resource generalist strain of *E. coli,* but not the loads of a specialized strain (*gat*-positive). (**a**) Analysis of the mouse gut microbiota divides mice into two main clusters, I (red) and II (blue). Dendrogram on the left represents UPGMA clustering of weighted Unifrac (beta diversity), with * indicating when jackknife node support was above 0.9 (100 subsamples of the OTU table). Bars on the right indicate the relative frequency of the different operational taxonomy units (OTUs; at 97% identity). Labeled taxa are enriched in either cluster I (red labels) or II (blue labels), as identified by LEfSe (full legend is provided in Fig. S4 and LefSe results in Fig. S5). (**b**) The gut microbiota affects the densities of *gat*-negative but not *gat*-positive *E. coli.* Dot plots indicate the densities of *E. coli* with each phenotype for the corresponding mouse-day combination. Line and shading indicate the mean and standard error. For *gat*-negative, two lines are shown as the bacterial densities differ significantly between red and blue clusters. Conversely, for *gat*-positive, only one line is shown as there is no significant difference. (**c-d**) Temporal dynamics of *gat*-negative and *gat*-positive *E. coli* showing that the densities of *gat*-positive clones remains constant across all mice, while the densities of *gat*-negative *E. coli* are lower in the mice with the cluster I microbiota (**c**), than in the mice with the cluster II microbiota (**d**). (**e**) Microbiota diversity (estimated as Shannon Index) is significantly lower in cluster II than in cluster I (similar observations are made for other diversity indices; see Fig. S6).

Importantly, cluster I is associated with an abundance of *gat*-negative *E. coli* that decreases overtime, tending to <10^8^ (per feces gram), which is significantly lower than that observed in cluster II (with average abundance of 10^9^ per feces gram; χ^2^_1_=33.6, *p*<0.001). These results strongly suggest that *gat*-negative *E. coli* competes with other members of the microbiota for resources and that the *gat*-positive *E. coli,* which acquired mutations leading to increase specialization on galactitol, may be selected for because it avoids such competition.

The findings reported here suggest that the gut ecology may favor natural polymorphism for galactitol metabolism in its microbiota. To test for this we sampled Enterobacteriacea clones from the natural microbiota of our host model organism (fecal samples of untreated mice in the IGC animal house (*n*=20)) and from the gut microbiota of healthy humans with no controlled diet (*n*=9). Remarkably, variation for this trait was found to be segregating both in mice and in man (Table S2). In 7 out of 9 human hosts polymorphism for ability to consume galactitol was detected and in 5 hosts the *gat*-negative phenotype was present at frequencies above 80%. From 20 mice assayed, in one the *gat*-negative phenotype was present at frequency, in another at low frequency and in the remainder only *gat*-positive clones were sampled. These results indicate that neither the antibiotic treatment nor the strain background used for the *E. coli* gut colonization, are the reason underlying the trait polymorphisms found here. Interestingly, in a survey of core genes in *E. coli, gatC* was found to be a target of repeated replacement substitution driven by positive selection (23).

Overall our results highlight how evolutionary and ecological processes interact to shape the genetic and phenotypic composition of commensal bacteria within a natural ecosystem. At the evolutionary level we find that a strong-mutation-strong-selection-strong-drift regime of adaptation can lead to the maintenance of polymorphism through frequent episodes of reverse evolution, which defy Dollo’s law of evolutionary irreversibility (24, 25). This regime of adaptation has been poorly studied, at both the theoretical and experimental levels, but maybe important for host-associated microbes as their population dynamics often involves transmission bottlenecks (strong-drift) followed by expansion to large population sizes (strong-mutation-strong-selection). Moreover, our study is pioneer at revealing that phenotypic and genetic reversals may be common in the mammalian intestine. Such scenario may have been difficult to predict, as bacteria could be expected to respond to changing environments by gene regulation and not through *de novo* mutation, as we shown here. One context where reversion is pivotal regards antibiotic resistance. Reversion of antibiotic resistant bacteria to sensitivity has been more rarely observed than what one would desire (26). However, since most studies have been done *in vitro*, it would be important to determine if the rate of reversion of resistance *in vivo* could be as high as the one we found here for a metabolic trait. At the ecological level, our results indicate that resource specialization may allow new emerging strains to avoid competition with other members of the microbiota (27), even if to stably colonize the gut at low abundances. Importantly, we could not have reached this conclusion solely from frequency data. Our results thus highlight the importance that absolute abundance data can have to understand ecological interactions among microbes. They further show how precise dietary supplementations may allow for controlling the colonization level of specific strains. In sum, the evolutionary invasion and ecological stability of *de novo* emerging clones in genetic identical hosts, nurtured in identical conditions, was highly reproducible, despite being accompanied by significant differences in the level of microbiota species richness between hosts.

## Materials and Methods

### Bacterial Strains and culture conditions

All strains used in this study were derived from MG1655, a K12 commensal strain of *Escherichia coli* (28).

*Ancestors of the first colonization:* the two ancestors of the first colonization (Fig. 3a), described in a previous experiment (5), were DM08-YFP and DM09-CFP (MG1655, *galK::YFP/CFP* amp^R^, str^R^ (*rpsL150), ΔlacIZYA)).*

*Ancestors of the second colonization:* strains JB19-YFP and JB18-CFP (MG1655, *galK::YFP/CFP* cm^R^, str^R^ (*rpsL150), ΔlacIZYA,* Ins (1bp) *gatC*) were used for the second colonization (Fig. 3b) and were previously described in (7). These strains differ from the ancestral MG1655 fluorescent strains DM08-YFP and DM09-CFP by a mutation in the *gatC* gene (1bp insertion in the coding region), rendering it auxotrophic for galactitol. To construct these strains the ampicillin resistance cassette in the ancestral strains DM08-YFP and DM09-CFP was replaced with a chloramphenicol resistant cassette using the Datsenko and Wanner method (29). The yellow (*yfp*) and cyan (*cfp*) fluorescent genes linked to cm^R^ were then transferred by P1 transduction to a derivative of clone 12YFP (6), an evolved clone of DM08-YFP, isolated after 24 days of adaptation in the gut of WT mice, that carried an insertion of 1bp in *gatC* and a large duplication. During the genetic manipulations the large duplication was lost, confirmed by whole genome sequencing, leaving this clone with a single mutation in *gatC* (7).

*Ancestor of the third colonization*: the strain used to start the third colonization was isolated from population 2.4 of the second colonization at day 24 post-colonization. This strain (named 19CFP (7) has a 2bp insertion (+TC) in *gatC* which reconstructed the open reading frame allowing it to recover the ability to consume galactitol (see below details about the phenotype confirmation). Additionally, this strain has the following mutations: an IS2 insertion in the coding region of *ykgB* and a 6790bp deletion from gene *intZ* to *eutA.*

Reconstruction of the transcriptional regulator of the galactitol metabolism (*gatR):* the *gatR* coding region is interrupted by an IS element in *E. coli* MG1655, and therefore in all strains used in this study. This insertion inactivates *gatR* leading to the constitutive expression of the *gat*-operon. To restore the function of the gene, and therefore the ability to regulate the operon, a copy of the *gatR* native sequence (obtained from the strain *E. coli* HS) was used to replace the non-functional copy. This was accomplished by the *SacB-TetA* counter selection method as previously described (30)). Briefly, the *SacB-TetA* fragment with the overhangs for recombination was obtained by target PCR using the primers *gatR_SacB_F /gatR_SacB_R* (Table S3) and genomic DNA of *E. coli* XTL298 as a template. The resulting PCR product was then used to replace the non-functional copy of *gatR* present in strains DM08 and DM09 by the Wanner method (29). The DNA fragment containing the native sequence of *gatR* was obtained by target PCR using the primers *yegS_F / gatD_R* and *E. coli* HS as a template. Finally, this fragment was used to replace the counter selection cassette in the *gatR* locus of the recipient strains DM08 and DM09 by the Wanner method (29) and plated on double-counter-selective medium. Candidates for successful restoration of *gatR* were screened by PCR and tested for antibiotic resistances to confirm the loss of the cassette and the plasmid before sequencing.

To distinguish between *gat*-negative and *gat*-positive bacteria we used the differential medium MacConkey agar supplemented with galactitol 1% and streptomycin (100μg/mL). Plates were incubated at 30°C. The frequency of *gat*-negative mutants was estimated by counting the number of white (auxotrophic for galactitol) and red colonies.

### *In vivo* Experimental Evolution

In order to study *E. coli’s* adaptation to the gut we used the classical streptomycin-treated mouse model of colonization (31) and performed the evolution experiments using the same conditions as before (5–7). Briefly, 6- to 8-week old C57BL/6 male mice raised in specific pathogen free conditions were given autoclaved drinking water containing streptomycin (5g/L) for one day. After 4 hours of starvation for water and food, the animals were gavaged with 100μL of a suspension of 10^8^ colony forming units of a mixture of YFP- and CFP-labelled bacteria (ratio 1:1) grown at 37°C in brain heart infusion medium to OD_600_ of 2. After the gavage, all animals were housed separately and both water with streptomycin and food were returned to them. Mice fecal samples were collected for 24 days and diluted in PBS, from which a sample was stored in 15% glycerol at −80°C and the remaining was plated in Luria Broth agar supplemented with streptomycin (LB plates). Plates were incubated overnight at 37°C after which fluorescent colonies were counted using a fluorescent stereoscope (SteREO Lumar, Carl Ziess) to assess the frequencies of CFP- and YFP-labelled bacteria. These fluorescent proteins are used as neutral markers with which we can follow to detect the emergence of beneficial mutations, since these markers hitchhike with the beneficial mutations that spread in the populations (8).

### *In vivo* competitive assays

To test the *in vivo* advantage of the evolving populations at the last time point of the evolution experiment (Fig. 1a) samples of either YFP or CFP clones isolated from day 24 (sub-populations) were competed against the respective ancestor labelled with the opposite fluorescent marker (n=2 for each population). These sub-populations were composed of mixtures of approximately 30 colonies with the same fluorescent marker isolated after plating the appropriate dilution of mice fecal pellets. The mixtures of clones were then grown in 10 mL of LB supplemented with chloramphenicol (100μg/mL) and streptomycin (100μg/mL) and stored in 15% glycerol at −80°C. *In vivo* competitions of evolved sub-populations against the ancestral were performed at a ratio of 1 to 1, following the same procedure described above for the evolution experiment. Mice fecal pellets were collected for 3 days, diluted in PBS and frozen in 15% glycerol at −80°C. Frequencies of CFP- and YFP-labelled bacteria were estimated using a fluorescent stereoscope (SteREO Lumar, Carl Ziess). The selective coefficient (fitness gain) of these mixtures of clones *in vivo* (presented in Fig. 1b) was estimated as: 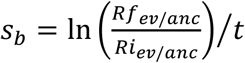, where *s_b_* is the selective advantage of the evolved clone, *Rf_ev/anc_ and Ri_ev/anc_* are the ratios of evolved to ancestral bacteria in the end (*f*) or in the beginning (*i*) of the competition and *t* is the number of generation per day. We assumed *t*=18 generations, in accordance with the 80 minute generation time estimated in previous studies on *E.coli* colonization of streptomycin-treated mouse (32, 33).

### Whole genome re-sequencing and mutation prediction

#### Clone analysis

Two *gat*-positive clones (revertant) isolated from the second colonization (day 6 of population 2.15) were WGS to test for the presence of additional mutations, besides the one reverting the *gat*-phenotype. Two independent colonies were used to inoculate 10 mL of LB and incubated at 37°C with agitation for DNA extraction (following a previously described protocol (34)). The DNA library construction and sequencing was carried out at the IGC sequencing facility. Each sample was paired-end sequenced on an Illumina Mi Seq Benchtop Sequencer. Standard procedures produced data sets of Illumina paired-end 250 bp read pairs. Genome sequencing data have been deposited in the NCBI Read Archive, http://www.ncbi.nlm.nih.gov/sra (accession no. SRP063701). Mutations were identified using the BRESEQ pipeline (35). To detect potential duplication events we used ssaha2 (36) with the paired end information. This is a stringent analysis that maps reads only to their unique match (with less than 3 mismatches) on the reference genome. Sequence coverage along the genome was assessed with a 250 bp window and corrected for GC% composition by normalizing by the mean coverage of regions with the same GC%. We then looked for regions with high differences (>1.4) in coverage. Large deletions were identified based on the absence of coverage. For additional verification of mutations predicted by BRESEQ, we also used the software IGV (version 2.1) (37).

#### Population analysis

DNA isolation was obtained in the same way as described above for the clone analysis except that now it derived from a mixture of >1000 clones per population grown in LB agar. Two populations, from the evolution experiment, were sequenced: 2.14 and 2.15. Those were sequenced for three time points during the adaptive period (generation 198 (day 11), generation 306 (day 17) and generation 432 (day 24)). The DNA library construction and sequencing was carried out by the IGC genomics facility. Each sample was pair-end sequenced on an Illumina MiSeq Benchtop Sequencer. Standard procedures produced data sets of Illumina paired-end 250 bp read pairs. The mean coverage per sample was between ∼100x and ∼300x for population 2.14 and between ∼70x and ∼115x for population 2.15. Mutations were identified using the BRESEQ pipeline (version 0.26) with the polymorphism option on. The default settings were used except for: a) requirement of a minimum coverage of 3 reads on each strand per polymorphism; b) eliminating polymorphism predictions occurring in homopolymers of length greater than 3; c) polymorphism predictions with significant (*P*=0.05) strand or base quality score bias were discarded. Data presented in Table S1.

### Diet supplementation experiment

To test the selective pressure exerted by galactitol, when present in the gut, we performed *in vivo* competitions between *gat*-positive (DM08) and *gat*-negative (JB18) bacteria, while supplementing the diet with this sugar. Competition experiments were performed as described above except that the initial frequency of *gat*-positive bacteria was ∼10% and the drinking water was supplemented with 0%, 0.01% or 0.1% of galactitol (n=3 for each galactitol concentration) together with streptomycin (5g/L). Fecal samples were daily collected and plated to access the frequencies of both phenotypes during four days.

### Estimate of gain-of-function mutation rate from *gat*-negative to *gat*-positive phenotype

Strains JB19 (*gatC* 1bp Ins) and 4YFP (*gatZ* IS Ins) were grown overnight in 10 mL of LB at 37°C with aeration. After growth, the total number of cells in the cultures was measured using BD LSR Fortessa (BD Biosciences) and approximately 1000 cells were used to inoculate 1 mL of LB (10 replicates of each strain) and incubated overnight. Aliquots of each replicate tube were plated in LB agar and MM agar supplemented galactitol 1% and incubated overnight at 37°C. The number of spontaneous *gat*-positive mutants and total number of cells grown on LB were used to estimate the mutation rate using the maximum likelihood approach as implemented in FALCOR (38).

We measured the mutation rate to re-acquire the ability to consume galactitol, since this locus was previously demonstrated to be a hotspot for loss of function mutations (7). The fluctuation test revealed that the spontaneous rate at which the ancestor of the second colonization (*gat*-negative) recovers the ability to consume galactitol is 9.7 x 10^−9^ (95% CI, [3 x 10^−9^, 1.9 x 10^−8^]) per generation. Considering that the estimated spontaneous rate of small indels in *E. coli* is ∼2x10^−11^ (39), and that about one third of such mutations in a region of ∼30 aminoacids of *gatC* could restore the open reading frame in the ancestral clone of the second colonization, the expected rate of mutation towards a *gat*-positive phenotype is ∼6x10^−10^. This estimate is between 5 to 15 times lower than that observed in the fluctuation assay, suggesting that a high mutation rate may have also contributed to the frequent re-emergence of *gat*-positive bacteria in the evolution experiment.

### Identification of adaptive mutations and estimate of haplotype frequencies in selected populations of the evolution experiment

In order to estimate the haplotype frequencies depicted in Fig. 2a-b two complementary strategies were employed. In addition to the WGS of the populations, targeted PCR of the identified parallel mutations was performed. For the targeted PCR, 20 to 80 clones from different time points were screened (from populations 2.14 and 2.15) using the same primers and PCR conditions as in (5, 7). Because all target mutations correspond to IS insertions an increase in size of the PCR band is indicative of the presence of an IS. Frequency of *gat*-revertant was estimated by plating in differential media (MacConkey supplemented with 1% galactitol). Frequencies are depicted in Table S1.

### Microbiota analysis

We extracted DNA from fecal samples of mice from the second colonization (Fig. 3b), at the 12 and 17 or 18 day post-colonization (*i.e*. 2 samples per mouse). Only mice where *gat*-positive *E. coli* could be detected where sampled (12 out of 15).

Fecal DNA was extracted with a QIAamp DNA Stool Mini Kit (Qiagen), according to the manufacturer’s instructions and with an additional step of mechanical disruption (40). 16S rRNA gene amplification and sequencing was carried out at the Gene Expression Unit from Instituto Gulbenkian de Ciência, following the service protocol. For each sample, the V4 region of the 16S rRNA gene was amplified in triplicate, using the primer pair F515/R806, under the following PCR cycling conditions: 94°C for 3 min., 35 cycles of 94°C for 60s, 50°C for 60s and 72°C for 105s, with an extension step of 72°C for 10min (41, 42). Samples were then pair-end sequenced on an Illumina MiSeq Benchtop Sequencer, following Illumina recommendations.

QIIME was used to analyze the 16S rRNA sequences through the following steps: i) sample de-multiplexing and quality filtering (using script split_libraries_fastq.py with default settings but with a phred quality threshold of 20); ii) OTU picking by assigning operational taxonomic units at 97% similarity against the Greengenes database (43); using script pick_open_reference_otus.py with default settings). This yielded variable sample coverage, with the lowest-coverage sample having 21735 reads. As the downstream analyses require even sampling, all samples were subsampled to 21735; iii) diversity analysis. The script core_diversity_analysis was used to obtain plots of relative abundance and estimates of alpha diversity (the different metrics shown in Fig. S6). As our dataset started with no pre-defined groups (i.e. the different mice did not correspond to particular treatment groups), UPGMA clustering of jackknifed UniFrac (weighted and unweighted) was used to identify microbiota clusters (44). This was done using the script jackknifed_beta_diversity by re-sampling 100 times at a depth of 16300 reads (75% of maximum read depth, as advised in the QIIME website) in order to obtain node support for each of nodes in the UPGMA tree.

As UPGMA clustering identified two main clusters (Fig. 4), linear discriminant analysis effect size (LefSe) was then used to identify if there were any taxa that were differentially abundant between the two clusters (45). The input data for LefSe (Fig. S8) was obtained after filtering the 97% OTUs for those that were present in at least 6 samples (*i.e.* at least 3 mice) and at a frequency that was above 0.5%, which corresponds to ∼100 reads.

### Natural fecal isolates collection and phenotyping

Fecal samples were collected from 9 mice of different litters (SPF-B6, strain C57BC-67) and from 9 healthy humans. Samples were weighted and re-suspended in PBS with 15% glycerol before storage at −80°C. To isolate lactose fermenting Enterobacteriaceae (red colonies) appropriate dilutions of the fecal samples were plated on MacConkey plates supplemented with 0.4% lactose and incubated overnight at 37°C. The frequency of *gat*-positive bacteria among the lactose fermenting Enterobacteriaceae was estimated by replica-plating ∼96 isolated clones on M9 minimal medium supplemented with 0,4% galactitol. The M9 agar plates were incubated at 30°C and bacterial growth was scored after 24h, 48h, 72h and 96h. The ability to metabolize galactitol was tested by three independent trials.

### Statistical Analysis

To analyze temporal dynamics data, we used linear mixed models, with mouse as a random effect. Where required, data were transformed to meet assumptions made by parametric statistics.

## Acknowledgments

We thank Karina Xavier, Luis Teixeira, Ivo Chelo, Armand Leroy, Christian Schlötterer, Michael Lassig and Jan Engelstaedter for discussions throughout this work. This research received funding from the European Research Council (ERC): ERC-StG-ECOADAPT; University of Cologne-Instituto Gulbenkian de Ciência, under SFB of DFG. The authors have no conflict of interest to report. The sequence data are available from http://www.ncbi.nlm.nih.gov/sra (accession no. SRP063701).

## Author Contributions

I.G. and A.S. designed the study with input from R. R. A.S., R.R., J.B., M. L., D.G. performed the experiments. A.S., R.R., M.L., I.G. analyzed the data and wrote the manuscript.

## Supporting Information

### Supplementary Text

#### General Model for Resource Competition

We make the very simplifying assumption that the gut constitutes a homogenous environment where bacteria compete for resources. We will use the model of competition for resources by Tilman (46) following van Opheusden *et al.* (19). We focus on the simplest model where we consider only two consumers and two resources in the mouse gut. The consumers are gat+ and *gat- E. coli,* whose densities inside the intestine are *Ec(t*) and En(t). The resources are galactitol and other general resources, whose abundances in the intestine are *G(t*) and *R(t*), respectively. We also assume that the system is homogeneous and that both strains of bacteria are removed at a similar rate (*d*). The rate at which bacteria grow (*f*) is assumed to solely dependent on the amount of resources available (*f* (*G,R*)). We further assume that, in the absence of bacteria, each resource achieves a maximum abundance *S_G_* and *S_R_,* for galactitol and other resources, respectively; and that the average residence time of either type of resource in the gut is 1/*a*. Resources are also depleted by each of the consumer strains. Under these assumptions the rate of change of each strain and each resource is:

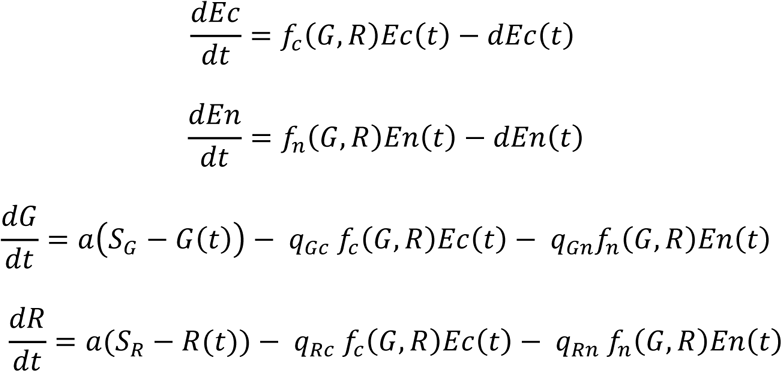

where *q_ji_* are the conversion factors for the resource *j* used by consumer *i*.

Under specific assumptions for the values of *q_ji_*, this system can be solved (19) such that the abundances of the two strains will tend to:

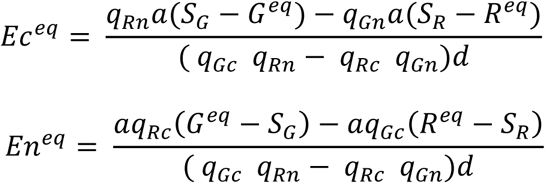

with the abundances of each resource dependent on the assumptions made for the relative growth rate of the strains, such that:

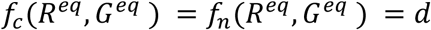

#### Model for colonization with gat+ and gat-strains

The *gat+* strain of *E. coli,* used in the first colonization, has an IS insertion in the repressor of the galactitol operon (*gatR*), which leads to the constitutive expression of this operon. Thus one can make the simplifying assumption that gat+ is specialized on galactitol such that its growth is only dependent on *G(t*); *i.e. f_c_ (G,R*)=*f_c_ (G*). On the other hand the *gat*-strain, which evolved from the *gat*+ during the colonization, does not consume galactitol, thus its growth is only dependent on *R(t*), *f_n_ (G,R)=f_n_ (R*). Under these assumptions the system simplifies to:

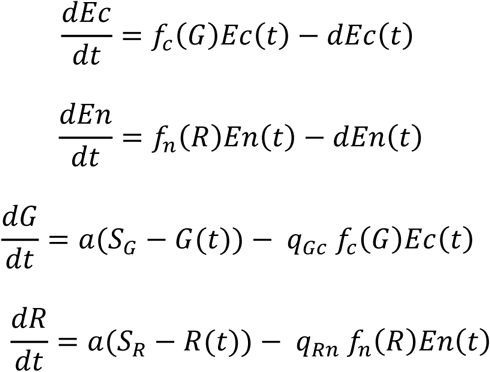

To move forward and calculate the equilibrium abundances of *gat*+ (*E_c_^eq^), gat- (E_n_^eq^*) and each of the resources *G^eq^* and *R^eq^*, we will assume a Michaelis-Menten dynamic for growth such that 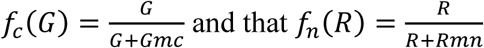, where *Gmc* and *Rmn* are constants. We then can deduce the equilibrium abundances of bacteria and resources to be:

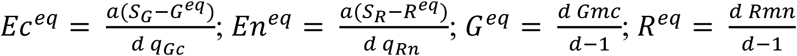

This simplified model implies that when *S_G_* increases, for example by supplementing the diet with galactitol, the loads of *gat*+ as well as the frequency of *gat*+ relative to *gat-*(which does not use this resource) should increase. Fig. S1a shows an example of the temporal dynamics for *gat*+ and *gat-* frequencies under this model. In Fig. S1b we show the prediction for the frequency of the strains under diet supplementation with increasing concentrations of galactitol.

#### Model for colonization with non-constitutive gat+ strain

We next asked whether a *gat*+ specialized strain, constitutively expressing the galactitol operon (*E_c_(t*)), could invade a generalist which can consume both galactitol and the other resources. *f_p_ (G,R*) is the growth function of such generalist, whose abundance is *E_p_(t*). If we assume that:

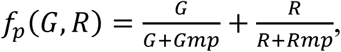

so that *E_p_* uses both resources, then we have:

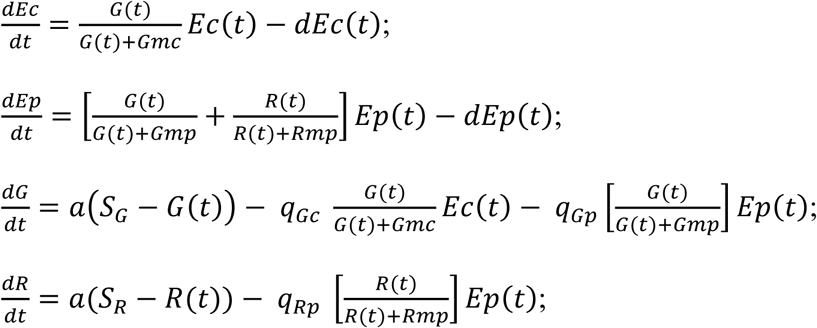

which leads to the expected abundances of:

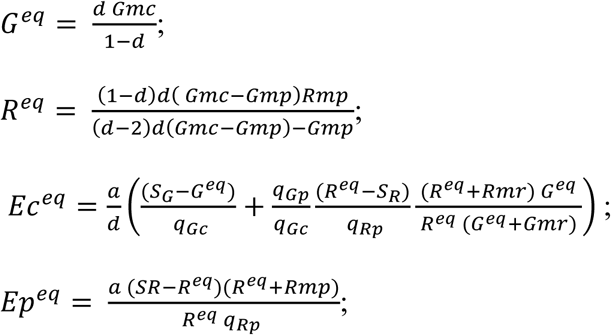

In Fig. S2 we show examples of parameter values where the emergence of a strain specialized on galactitol can rise in frequency.

**Fig. S1.**
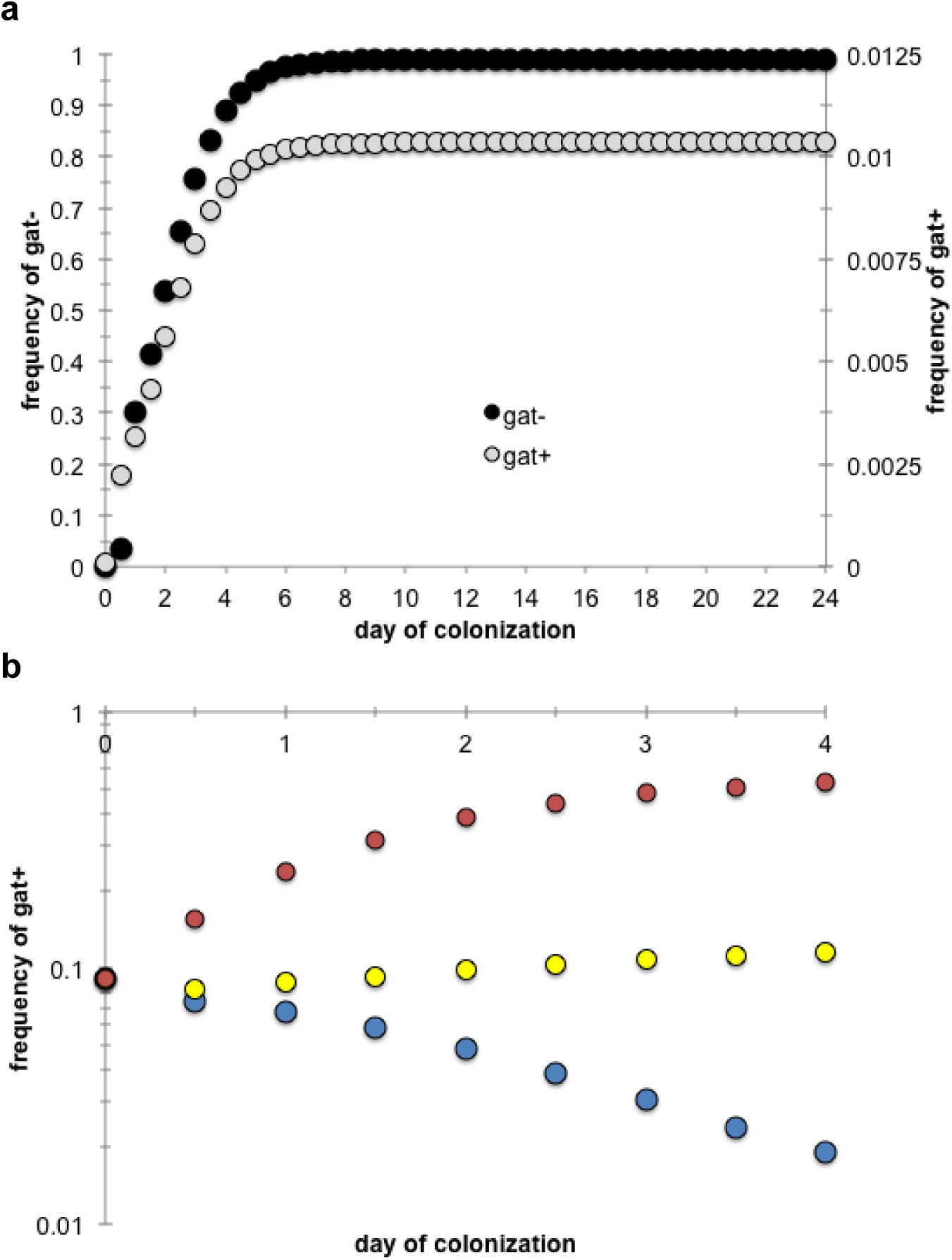
Dynamics of frequency change of *gat+* and *gat-* clones under the two-strain two-resource model where *gat+* only consumes galactitol. Model parameters are as follows: *S_R_*=0.02; *S_G_*=0.002; *a*=*d*=0.042; *Gmc*= 2x10^−5^; *q_Gc_*= 10^−3^; Rmn=2x10^−2^; *q_Rn_* =10^−4^. (**a**) Black circles: dynamics of *gat-* emergence in a mono colonization with *gat+;* gray circles: dynamics of *gat+* emergence in a mono colonization with *gat-;* (**b**) Dynamics of *gat+* frequency (as log_10_) under diet supplementation with: *S_G_*=0.002 in blue; *S_G_*=0.027 in yellow; and *S_G_*=0.252 in red.

**Fig. S2.**
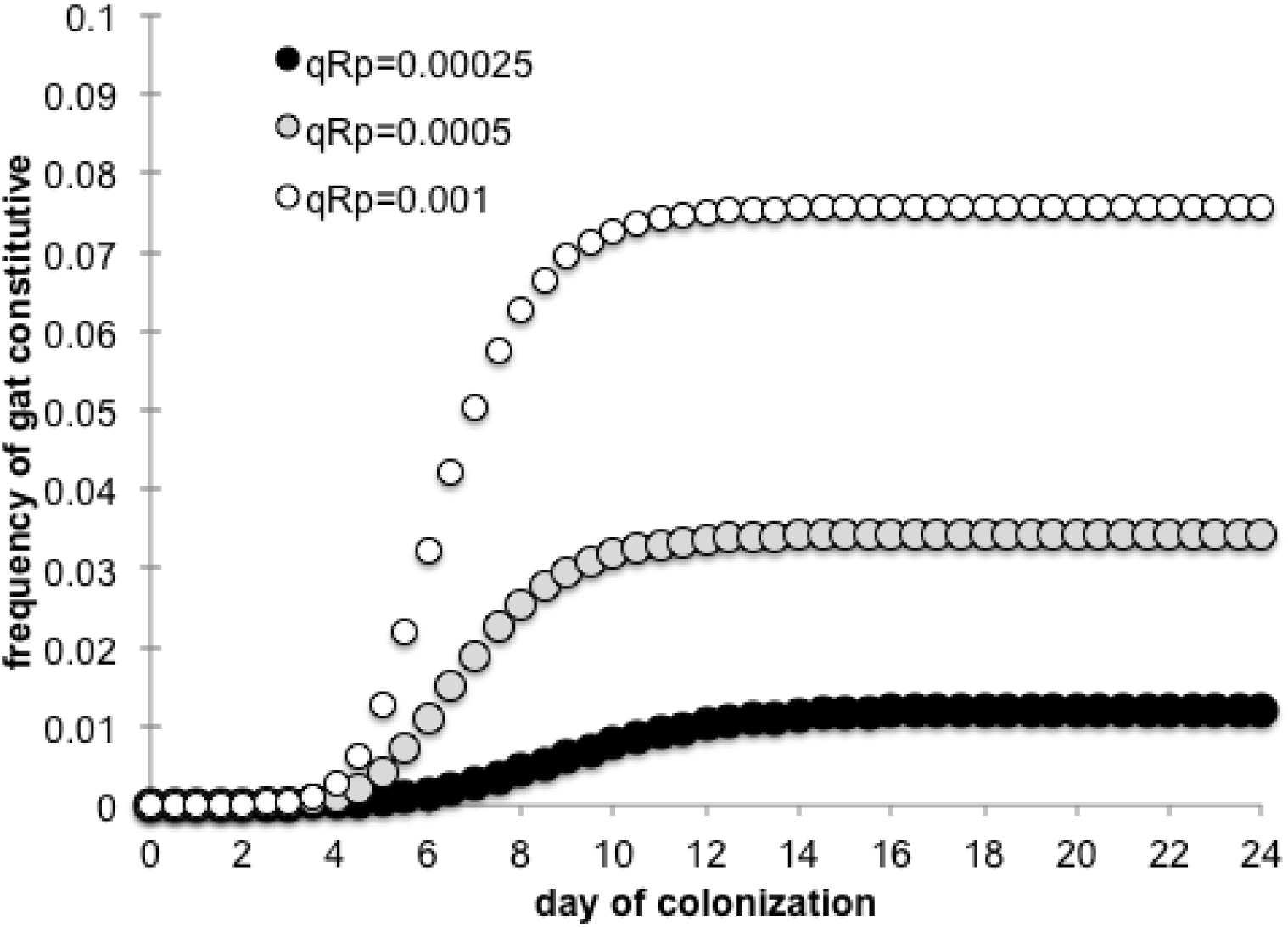
Dynamics of frequency change of the gat+ strain, which constitutively expresses the *gat* operon and thus is assumed to only consume galactitol, when invading a resident population of gat+ clones, which can consume both galactitol and the other resource. Model parameters are as follows: *S_R_*=0.02; *S_G_*=0.002; *a*=*d*=0.042; *Gmc*= 2x10^−5^; *q_Gc_*=10^−3^; *Rmp*=2x10^−2^; *Gmp* = 2x10^−4^; *q_Gp_*= 10^−4^; and *q_Rp_* as indicated.

**Fig. S3.**
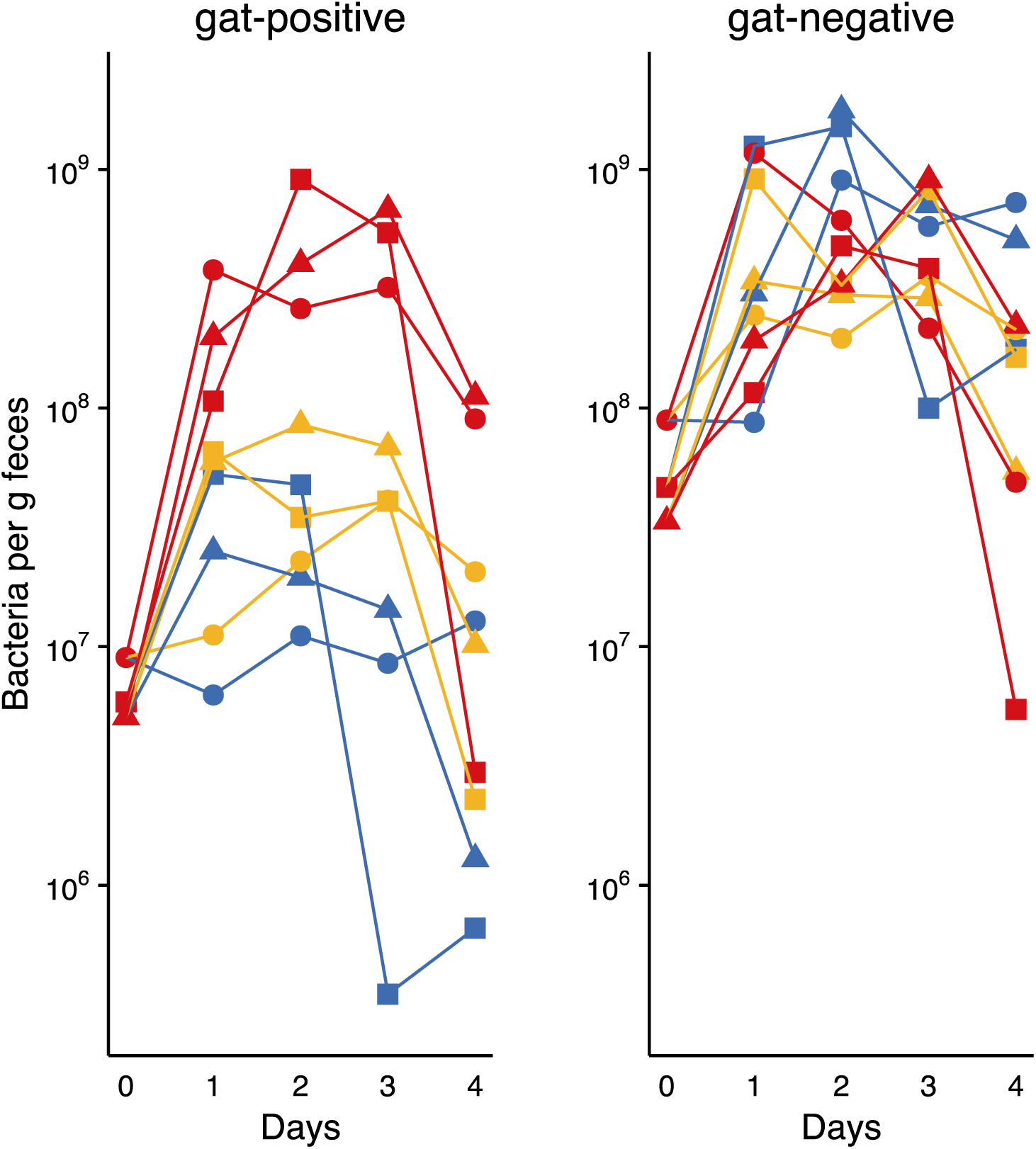
Temporal dynamics for the density of *gat*-positive (left) and *gat*-negative (right) bacteria for mice whose diet was supplemented with different galactitol concentrations. Colors correspond to those in Fig. 3d: blue – no galactitol supplementation; yellow – 0.01% galactitol; red – 0.1% galactitol (galactitol was diluted in the mice drinking water). Different symbols correspond to different mice.

**Fig. S4.**
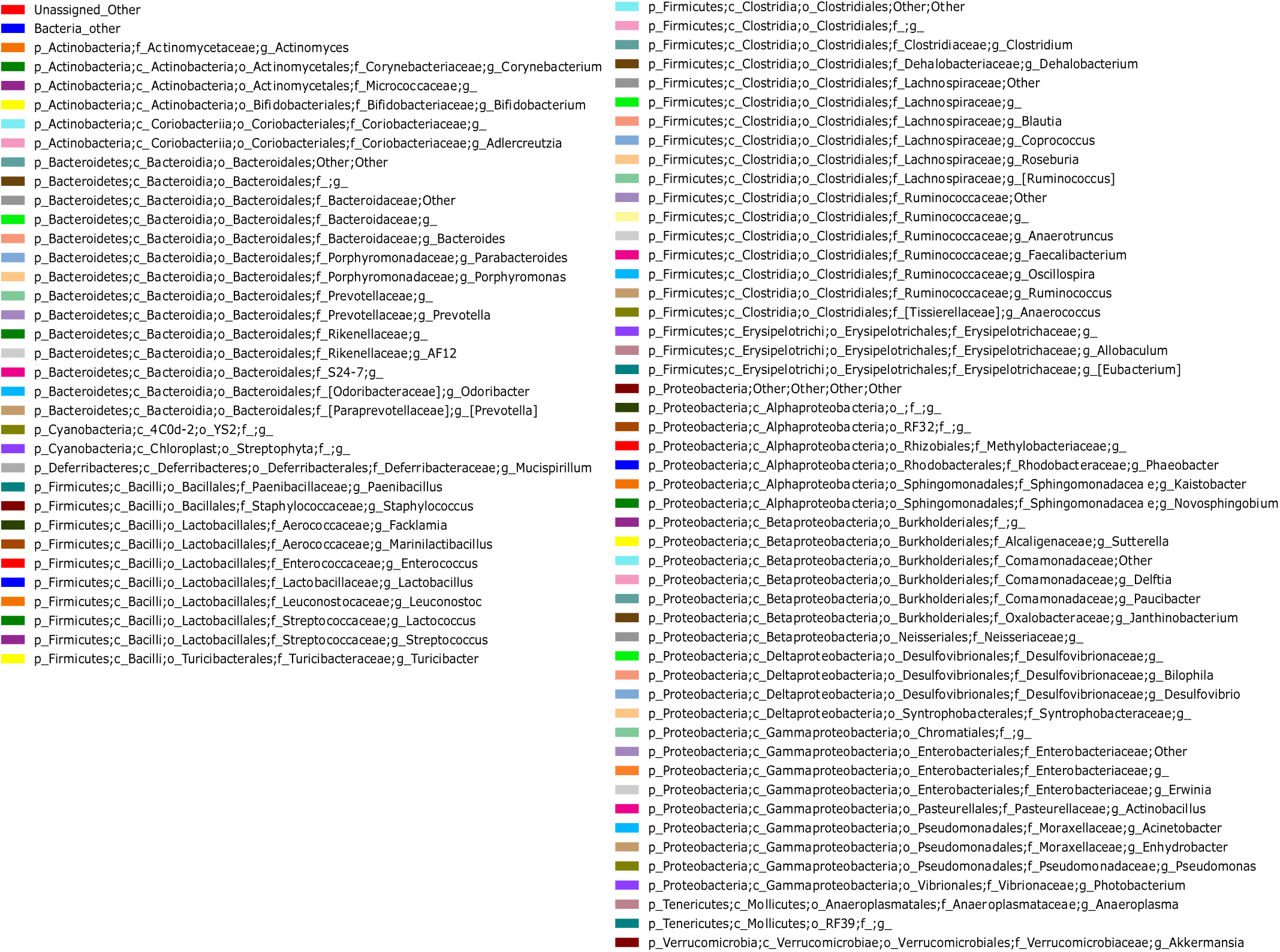
Full legend for Fig. 4a.

**Fig. S5.**
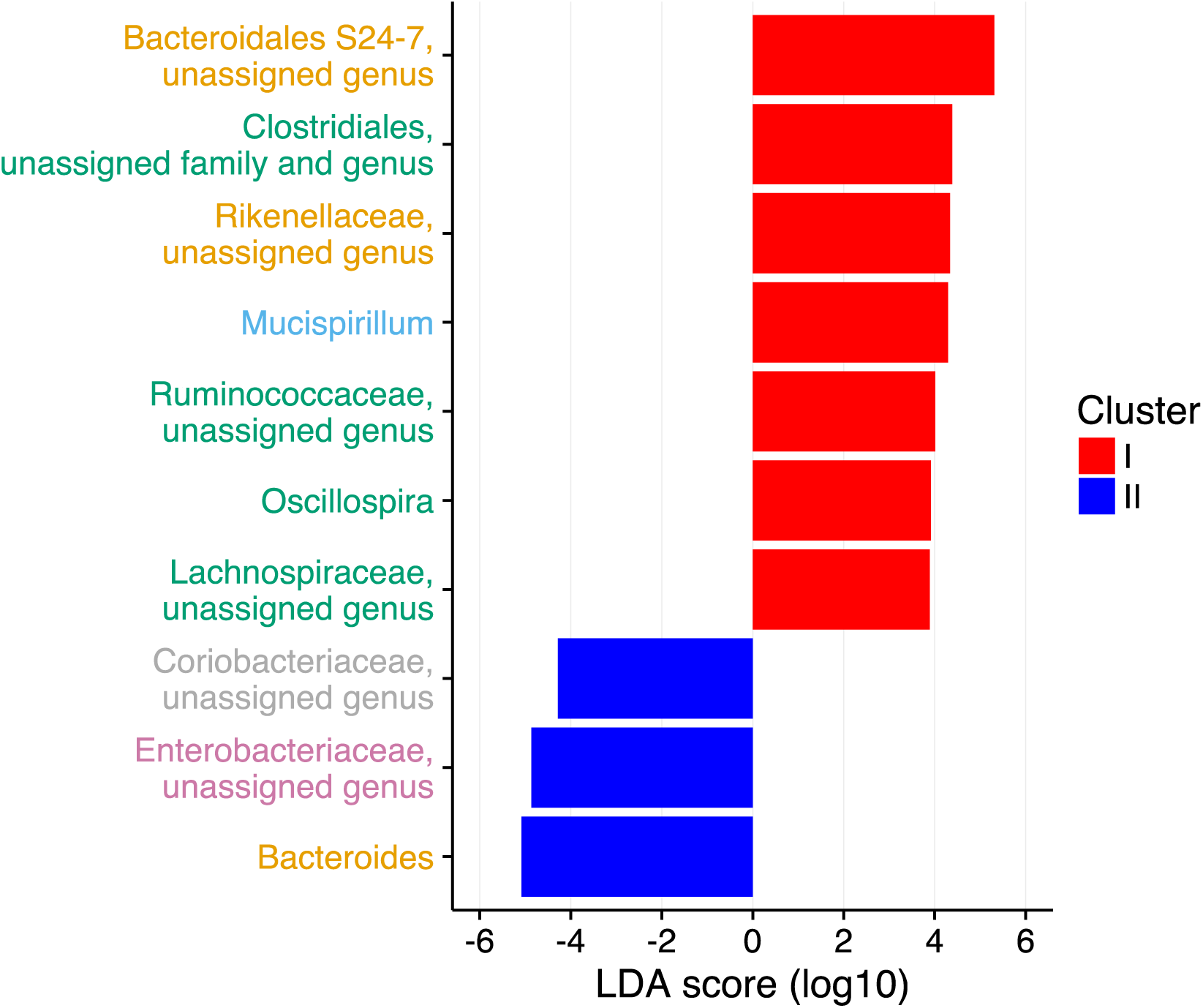
Linear discriminant analysis score (LDA) of 97% OTUs enriched for either cluster I (red) or cluster II (blue), as inferred using LefSe. OTUs are coloured according to their phylum: Actinobacteria (grey), Bacteroidetes (orange), Deferribacteres (blue), Firmicutes (green) and Proteobacteria (purple).

**Fig. S6.**
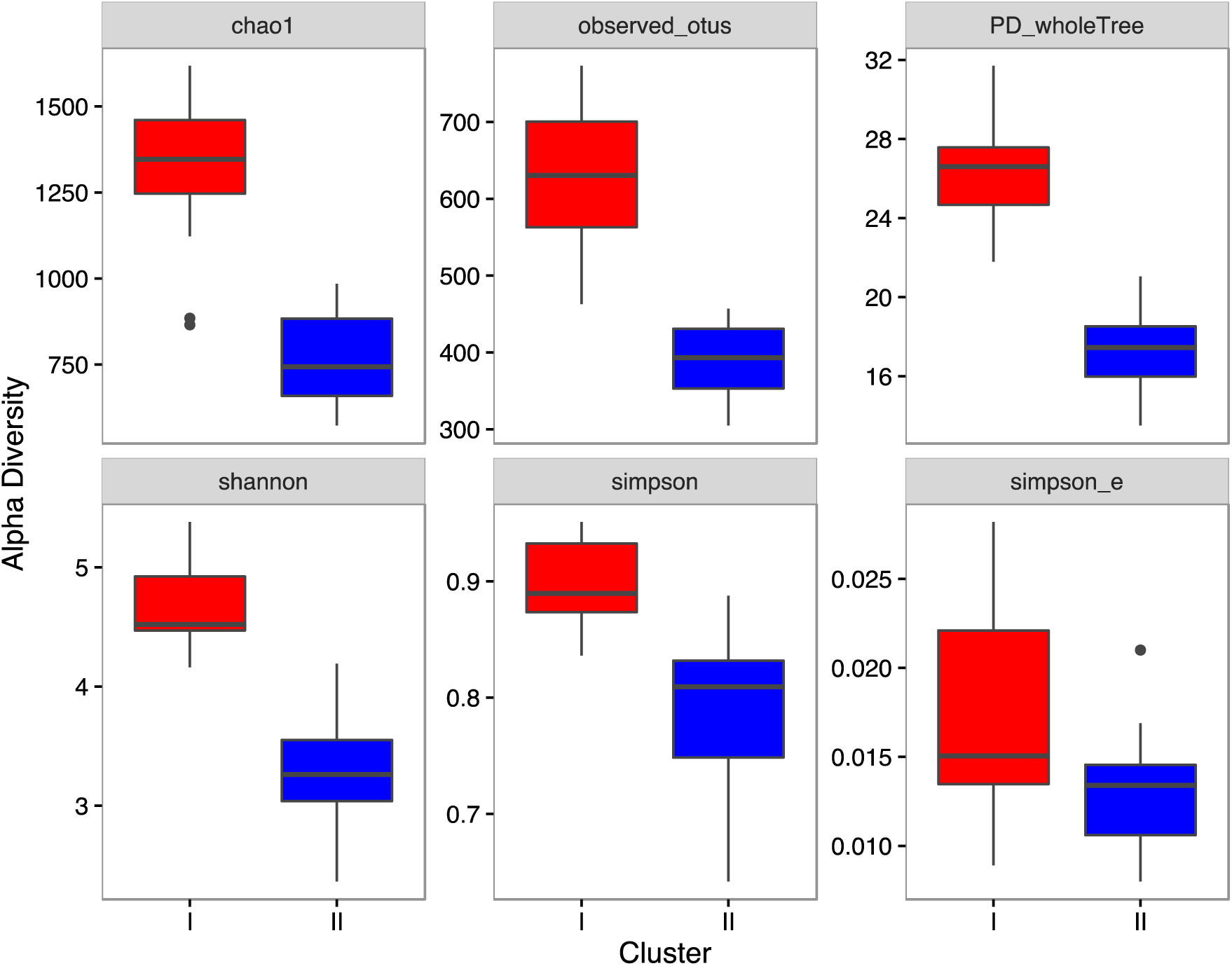
Alpha diversity is higher in cluster I than in cluster II, across multiple indices. chao1 - chao1 richness estimator; observed_otus – number of distinct OTUs; PD_wholeTree – Faith’s phylogenetic diversity; shannon – Shannon entropy; simpson – Simpson index; simpson_e – Simpson’s evenness. Significant differences between the two clusters (p<0.05, determined from linear mixed effects models) are detected for all indices except Simpson’s evenness.

**Table S1.**
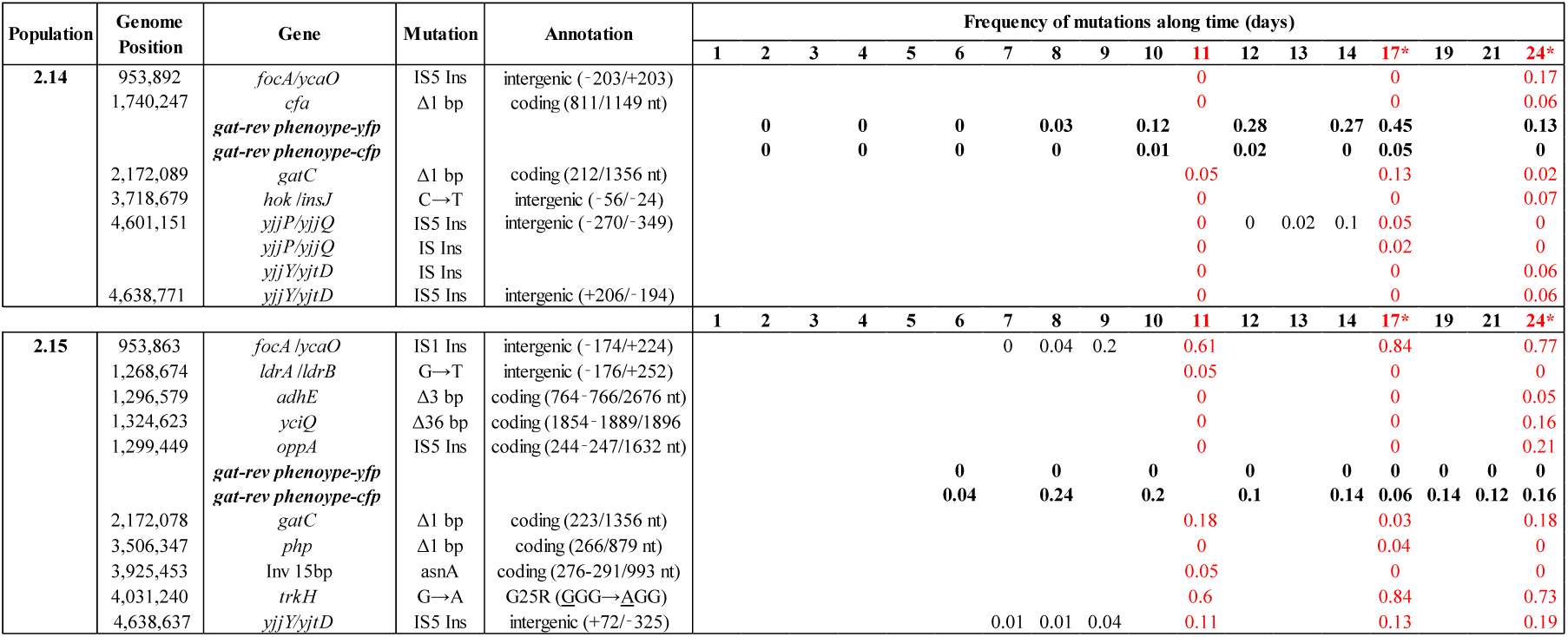
Genetic and phenotypic (in bold) polymorphism in two evolved populations. Mutations were detected by whole genome sequencing of samples of each population at different time points (shown in red) or by target sequencing (shown in black). In addition the frequency of the *gat*-rev phenotype in each of the fluorescence backgrounds was estimated by plating in selective media (shown in bold). The full data is graphically represented in Fig. 2.

**Table S2.** Polymorphism for galactitol metabolism in natural fecal isolates.

| **Sample #** | ***gat*-pos** | ***gat*-neg** | **Total** | **Frequency**<br>**of *gat*-neg** | **2SE** |
| --- | --- | --- | --- | --- | --- |
| Mouse 1 | 0 | 48 | 48 | 1 | 0.02 |
| Mouse 2 | 42 | 6 | 48 | 0.125 | 0.09 |
| Mouse 3-20 | 48 | 0 | 48 | 0 | 0.02 |
| Human 1 | 95 | 1 | 96 | 0.01 | 0.02 |
| Human 2 | 12 | 84 | 96 | 0.88 | 0.07 |
| Human 3 | 96 | 0 | 96 | 0 | 0.01 |
| Human 4 | 1 | 95 | 96 | 0.99 | 0.02 |
| Human 5 | 10 | 86 | 96 | 0.90 | 0.06 |
| Human 6 | 48 | 48 | 96 | 0.50 | 0.10 |
| Human 7 | 1 | 95 | 96 | 0.99 | 0.02 |
| Human 8 | 10 | 86 | 96 | 0.90 | 0.06 |
| Human 9 | 95 | 0 | 95 | 0 | 0.01 |

**Table S3.**
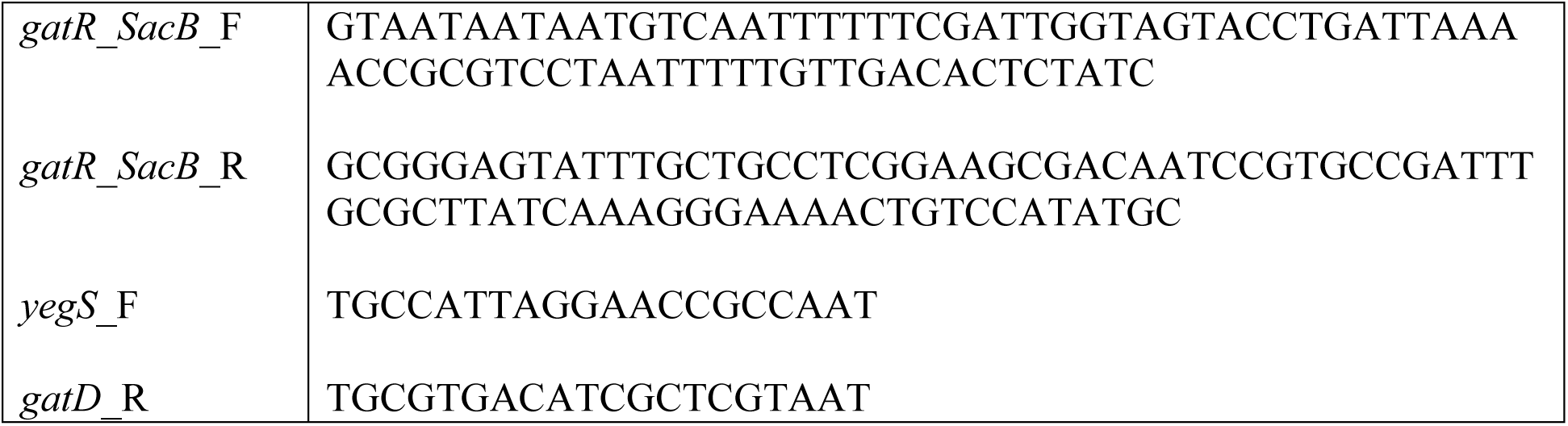
Primers used for the reconstruction of the transcriptional regulator of the galactitol metabolism (*gatR*)

